# Precise spatial representations in the hippocampus of a food‑caching bird

**DOI:** 10.1101/2020.11.27.399444

**Authors:** Hannah L. Payne, Galen F. Lynch, Dmitriy Aronov

## Abstract

The hippocampus is an ancient neural circuit required for the formation of episodic memories. In mammals, this ability is thought to depend on well-documented patterns of neural activity, including place cells and sharp wave ripples. Notably, neither pattern has been found in non-mammals, despite compelling examples of episodic-like memory across a wide range of vertebrates. Does episodic memory nonetheless have a universal implementation across distant neural systems? We addressed this question by recording neural activity in the hippocampus of the tufted titmouse – an intense memory specialist from a food-caching family of birds. These birds cache large numbers of food items at scattered, concealed locations and use hippocampus-dependent memory to retrieve their caches. We found remarkably precise spatial representations akin to classic place cells, as well as sharp wave ripples, in the titmouse hippocampus. These patterns were organized along similar anatomical axes to those found in mammals. In contrast, spatial coding was weaker in a different, non-food-caching bird species. Our findings suggest a striking conservation of hippocampal mechanisms across distant vertebrates, in spite of vastly divergent anatomy and cytoarchitecture. At the same time, these results demonstrate that the exact implementation of such common mechanisms may conform to the unique ethological needs of different species.

## Introduction

The hippocampus is critical for forming memories of experienced events, or episodic memories^1,2^. It has been a challenge to relate this function to specific patterns of neural activity. A central, experimentally tractable component of episodic memories is the storage of spatial information about *where* an experience occurred. In the mammalian hippocampus, spatial memory is thought to depend on two striking features of neural activity. During movement, place cells precisely represent spatial location^3^, whereas during wakeful immobility or sleep, firing patterns are reactivated in synchronized events called sharp wave ripples (SWRs)^4^. It is hypothesized that place cells represent the spatial component of episodic memories^5^, whereas SWRs reactivate and consolidate the memories^6,7^.

An enduring mystery is that place cells and SWRs have not been identified in the hippocampus of non-mammals, including multiple bird species, despite numerous recordings^8–13^. In spite of this absence, compelling examples of episodic-like, spatial memory have been demonstrated in a wide range of species, including fish, birds, and non-avian reptiles^^14–16^^. Memory in these non-mammals relies on a hippocampal structure^17,17,18^ that shares embryological origins with its mammalian counterpart^19,20^ and expresses many of the same genetic markers^19–21^. Given the apparent differences in hippocampal activity between species, it is therefore unclear which features of neural activity are fundamental, and which are dispensable, for hippocampus-dependent spatial memory.

The reported lack of mammalian-like hippocampal activity in non-mammals may not be surprising given dramatic differences in hippocampal anatomy across species. In mammals, sensory inputs indirectly reach the hippocampus after extensive processing by a six-layered cortex^22,23^. Inputs from different cortical regions are anatomically organized and are thought to account for functional specializations of different parts of the hippocampus^24^. The hippocampus itself is laminated and features a prominent pyramidal cell layer. In contrast, non-mammals have neither a six-layered cortex^25^ nor a hippocampal pyramidal cell layer^26,27^, and hippocampal inputs are not known to be organized. These differences raise the question of how similar hippocampal memory functions are implemented by fundamentally different neural circuits across species.

To address this question, it is critical to examine neural activity across species that have comparable mnemonic abilities. We therefore focused on a non-mammal with exceptional spatial memory: a food-caching bird. Food-caching birds store many individual pieces of food at scattered, concealed locations in their environment and later retrieve these caches using memory^15,16^. Accurate retrieval requires the hippocampus, which is grossly enlarged in these memory specialists^17,18,28^. To investigate what physiological and anatomical substrates might enable robust spatial memory, we systematically recorded throughout the avian hippocampus, both during locomotion and during sleep.

## Results

We first asked whether spatial activity could be detected in the hippocampus of a food-caching bird. We designed miniature microdrives that allowed unconstrained locomotion of a small bird in a two-dimensional arena. These microdrives were implanted into the hippocampus of tufted titmice — food-caching members of the *Paridae* family, which also includes chickadees^29^ (Extended Data Fig. 1). For comparison with classic studies of the rodent hippocampus, we recorded single cells while titmice foraged for sunflower seed fragments randomly dispensed throughout the arena (**Fig. 1a-c** and Extended Data Fig. 2).

**Figure 1:**
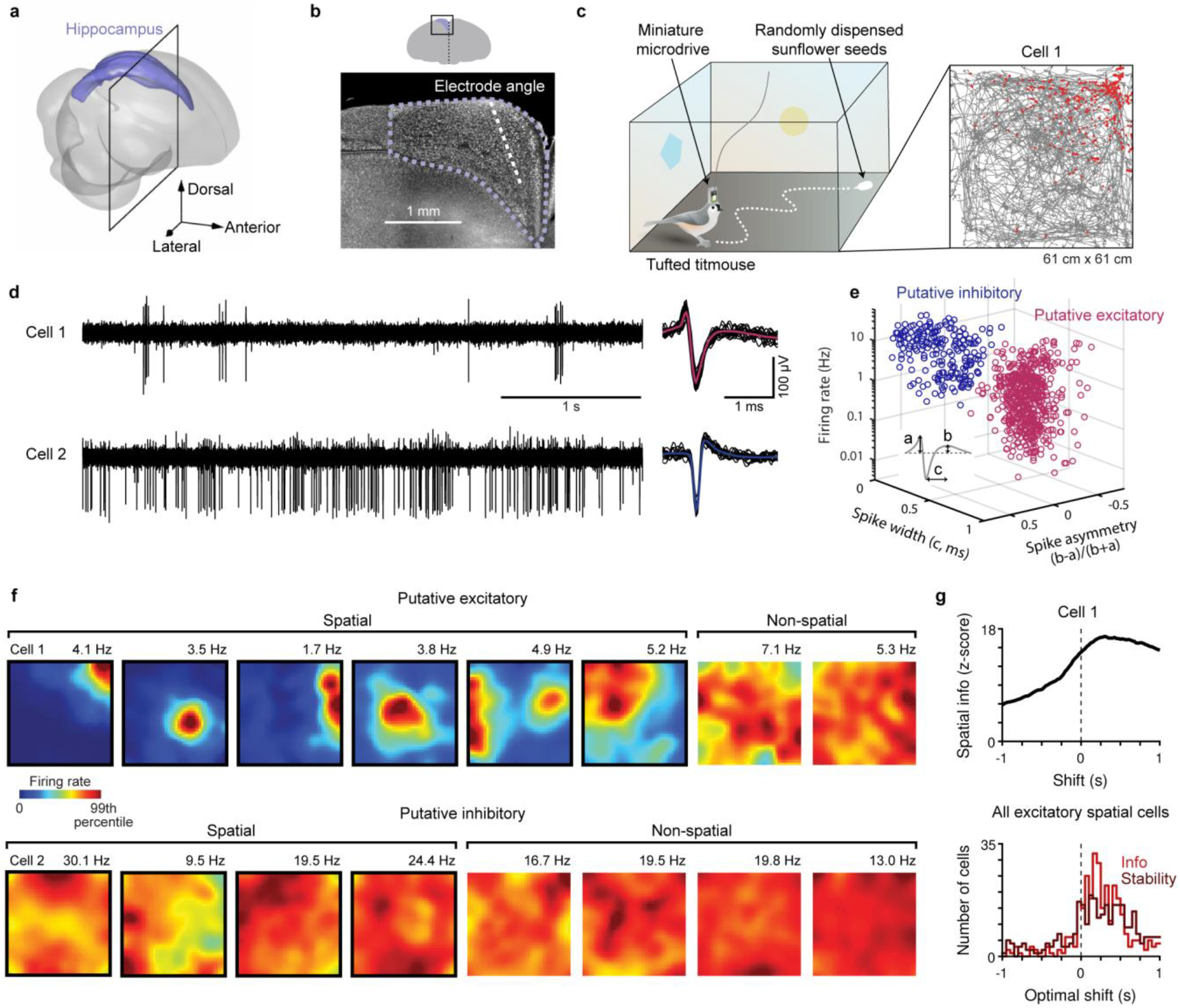
Place cells in the hippocampus of tufted titmice. **a**, Reconstruction of the titmouse hippocampus. **b**, Fluorescent Nissl-stained coronal section at the location indicated by the black box in **a**. Dashed purple: hippocampal boundary. Dashed white: electrode approach angle. **c**, Left, schematic of the random foraging arena. Right, bird’s trajectory (grey line) and locations of spikes (red dots) for an example hippocampal cell. Cell 1 refers to the same neuron in all panels. **d**, Voltage traces and 20 example spike waveforms for two example cells (black: examples; pink or blue: mean). **e**, Electrophysiological characteristics for all cells recorded during the random foraging task, classified as excitatory (n = 538) and inhibitory cells (n = 217) (see Methods). **f**, Example spatial rate maps for excitatory and inhibitory neurons. Numbers above plot indicate maximum of color scale. **g**, Top, spatial information as the time shift between spikes and behavior was varied for an example cell. The peak at a positive shift (“optimal shift”) means that spikes were most informative about the bird’s future position. Bottom, histogram of optimal shifts for spatial information and spatial stability.

**Figure 2:**
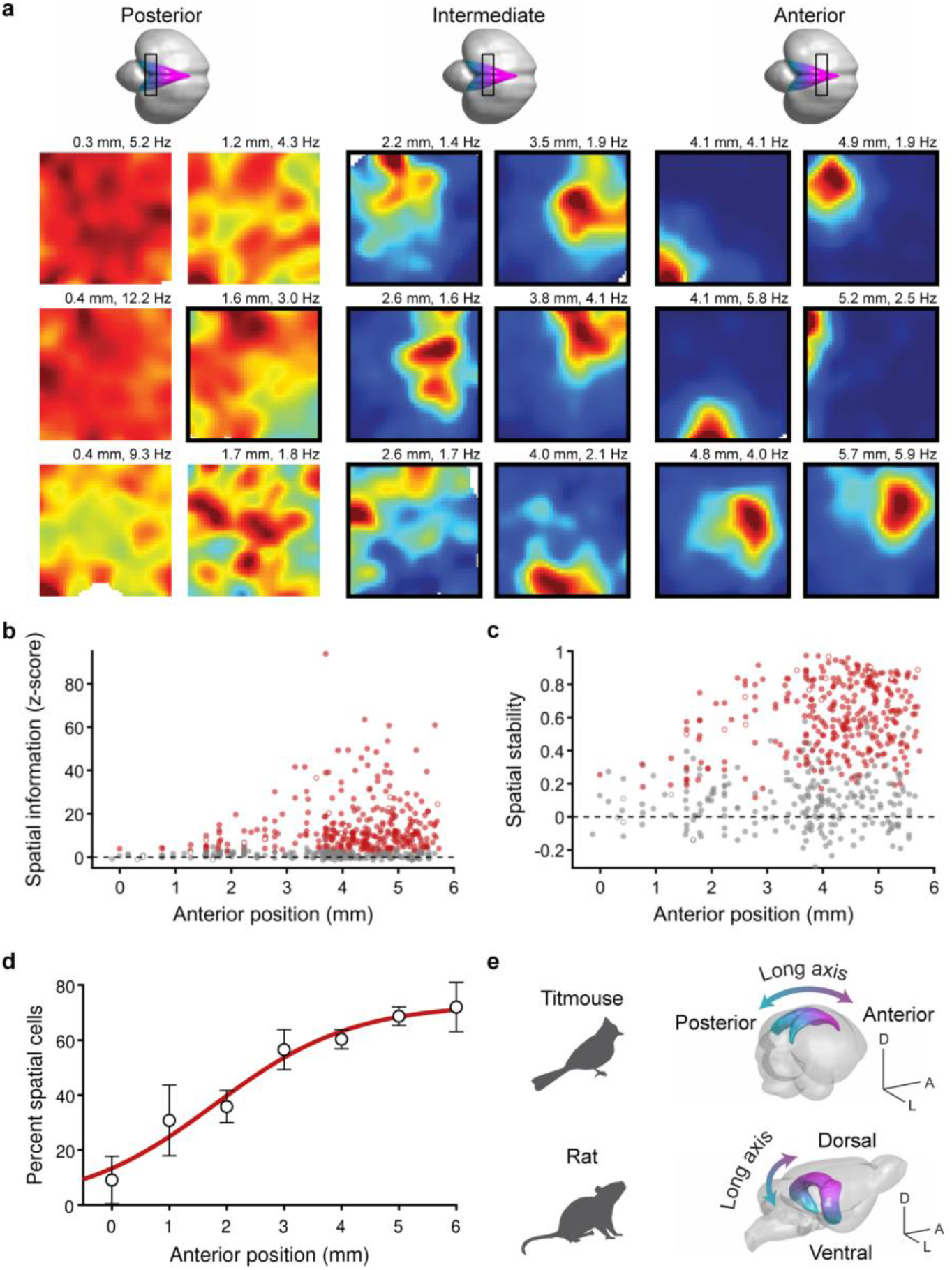
Spatial representations are organized across the long axis of the hippocampus. **a**, Example spatial rate maps for excitatory neurons from posterior, intermediate, or anterior hippocampus, plotted as in **Fig. 1**. Place cells are outlined in black. The location on the anterior-posterior axis (distance from lambda) is indicated above each map. **b**, Spatial information, normalized by taking the z-score of the actual value relative to a shuffled dataset, plotted for all 538 excitatory cells. Red: place cells; grey: non-place cells; open markers: example cells in **a**. **c**, Spatial stability plotted as in **b**. **d**, Fraction of excitatory cells that passed place cell criteria binned across anterior position. Error bars: mean ± SEM; red line: logistic sigmoid function fit. **e**, Schematic of the spatial gradient across the hippocampal long axis in tufted titmice and in rats (3D model generated using published data^50^). Scale bars 5 mm.

Birds are far removed from mammals in evolution^30^, and it is unknown whether the basic cellular components of the hippocampus are comparable across species. As a preliminary step, we therefore analyzed electrophysiological characteristics of the recorded neurons. This analysis revealed two prominent clusters (n = 538 and 217 cells). Cells in the first cluster had lower firing rates, wider spikes, and a larger first peak of the spike waveform than cells in the second cluster (**Fig. 1d,e**). Cells in the first cluster were also more likely to fire in bursts (CV2 1.14 (SD 0.15) and 0.86 (SD 0.14), respectively, p = 10^-88^, two-sided t-test). Together, these characteristics are a remarkable match to those of excitatory and inhibitory neurons in the mammalian hippocampus, respectively^31^. In fact, we occasionally recorded pairs of cells on the same electrode, and in a large fraction of these cases, the cross-correlogram of spike times suggested a monosynaptic connection consistent with the putative identities (Extended Data Fig. 3). The basic classes of hippocampal neurons therefore appear to be conserved between birds and mammals. We will refer to cells in these classes as excitatory and inhibitory neurons.

**Figure 3:**
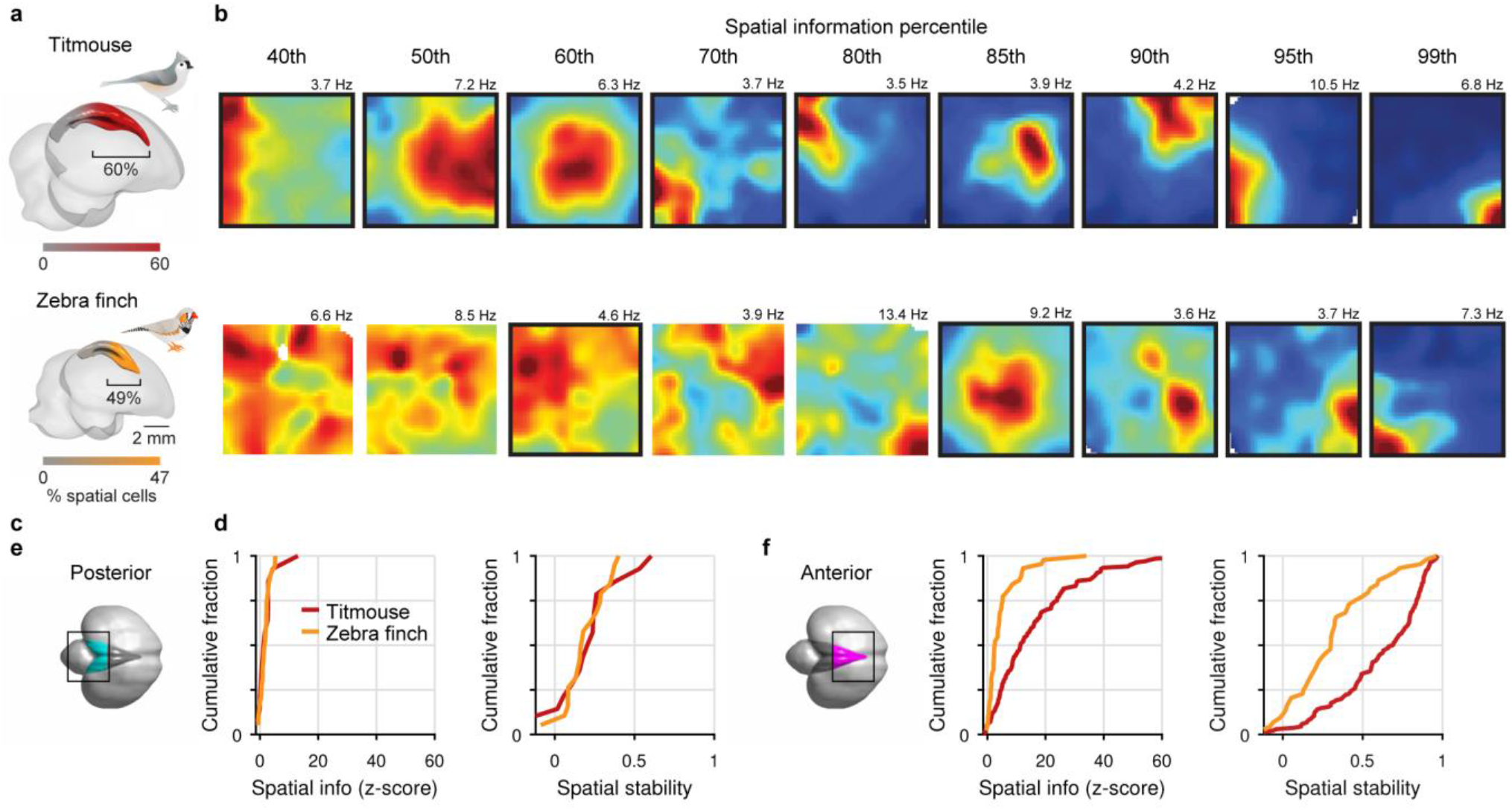
Spatial representations differ across avian species. **a**, Titmouse (top) and zebra finch (bottom) hippocampus colored according to a logistic sigmoid fit to the percent of place cells at each anterior position. The bracket indicates the percent of hippocampal length anterior to the inflection point of this fit. **b**, Example spatial rate maps. All cells within the bracketed region in **a** with peak rates >3 Hz were ranked by spatial information, and rate maps for the cells corresponding to the given percentiles are shown. Place cells are outlined in black. **c**, Cumulative distributions of normalized spatial information and spatial stability for cells from the posterior hippocampus, as defined anatomically (black box; see Methods; n = 14 and 19 for titmouse and zebra finch, respectively). **d**, Same as **c** but for anterior hippocampus (n = 136 and 44, respectively).

Surprisingly, the firing of many neurons in the titmouse hippocampus was precisely localized in space (**Fig. 1f**). We used two conventional criteria to identify such cells. First, we measured “spatial information” as the mutual information between firing rate and location. Second, we measured “spatial stability” by cross-correlating firing rate maps from disjoint subsets of the recording session. Many neurons passed criteria for both: significantly high spatial information and significantly high spatial stability (quantified below). Of these cells, excitatory neurons often displayed spatially restricted firing fields that were highly reminiscent of rodent place fields. Across the population, significant firing fields fully tiled the environment (Extended Data Fig. 4). In rats, place cells are not simply responsive to the animal’s location, but predict future position by 100-200 ms^32^. Despite different methods of locomotion in titmice and rats (discrete hops vs. continuous walking), we found that titmouse hippocampal cells were also most strongly tuned to future position (mean best delay 220 ms for spatial information and 204 ms for spatial stability; both greater than 0, p < 10^-14^, two-sided t-test, n = 309 and 307, respectively; **Fig. 1g**). Additionally, some neurons displayed head direction and speed tuning (Extended Data Fig. 5). In separate experiments, neurons also displayed directional tuning on a linear track (Extended Data Fig. 6). The titmouse hippocampus therefore displays many features of activity previously observed in mammals and, notably, contains neurons resembling classic place cells. We will proceed to use the term “place cell” for excitatory neurons with significantly high spatial information and stability.

**Figure 4:**
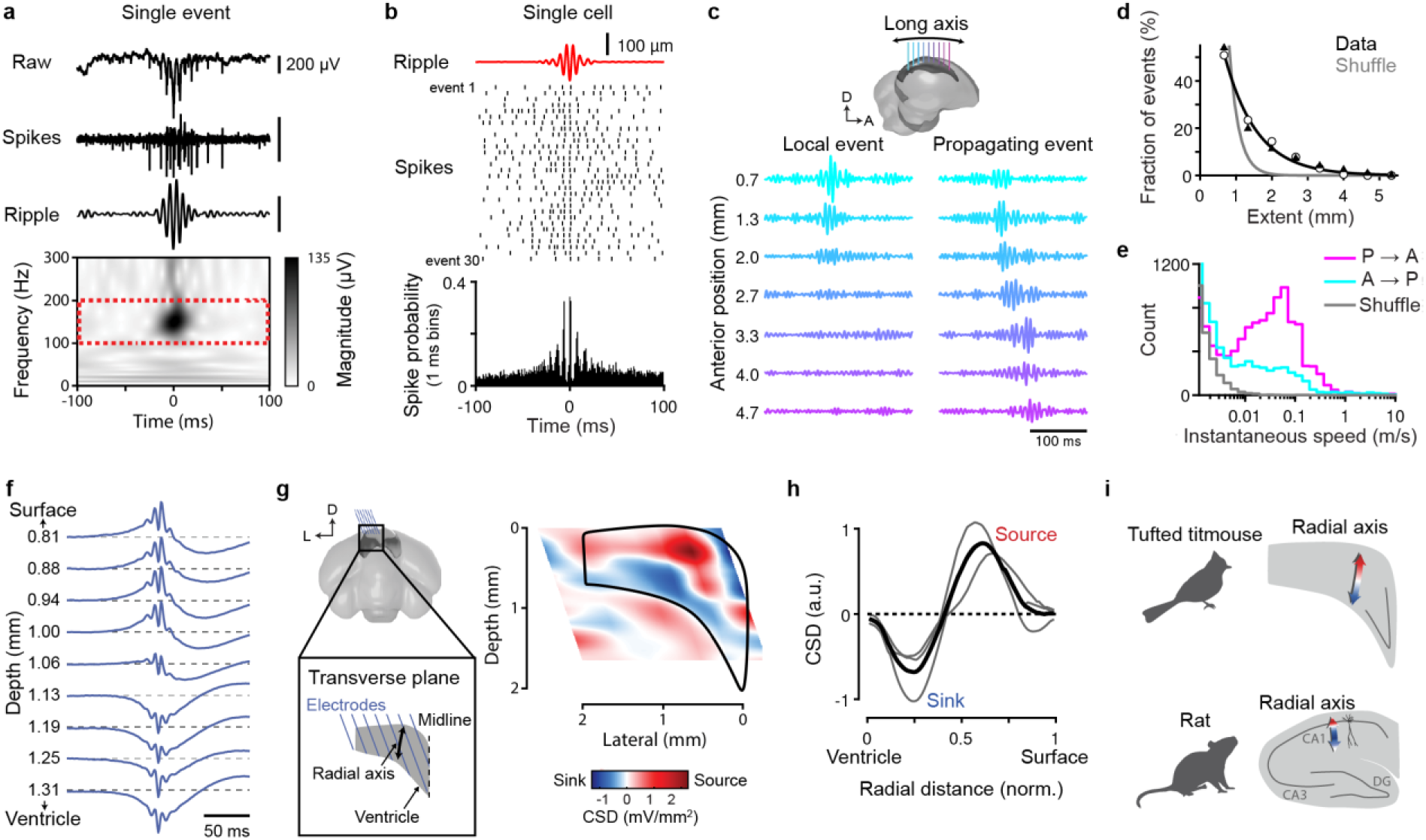
SWRs in the avian hippocampus. **a**, Single SWR in the titmouse hippocampus. Top: raw, spike-band filtered (800–5000 Hz), and ripple-band filtered (100–200 Hz) LFP. Bottom: spectrogram of the LFP. **b**, Average ripple-band LFP, spike raster from 30 consecutive SWRs, and spike histogram across all detected SWRs for a single cell. **c**, Top: schematic of recording configuration across the long axis of the hippocampus. Bottom: two individual SWRs detected on multiple electrodes. The “local” event is restricted to a small number of electrodes, whereas the “propagating” event extends over 4 mm. **d**, Distribution of the extent of SWRs across the long axis. Markers: two individual titmice; black line: exponential fit to the mean of both birds; grey line: exponential fit to shuffled data. **e**, Speed of SWRs propagating in the posterior-to-anterior (P→A) and anterior-to-posterior (A→P) directions, compared to speed for a shuffled data set (see Methods). **f**, Raw LFP averaged across SWRs for a single electrode at different depths in the titmouse hippocampus. **g**, Left: schematic of electrode placement within the transverse plane of the hippocampus. Right, 2D CSD map within the transverse plane of the hippocampus (black outline) in one bird. **h**, One-dimensional CSD collapsed across the radial axis for three titmice (grey lines: individual birds; black: average). Normalized radial distance is 0 at the ventricle and 1 at the dorsal surface of the hippocampus. **i**, Layered CSD organization during SWRs in titmouse and in rat^37^. In rat, the primary current sources (red) and sinks (blue) correspond to the pyramidal cell layer and the stratum radiatum, respectively. Note that rat CA1 is folded upside down relative to the avian hippocampus in stereotaxic coordinates. As a result, sources and sinks are actually reversed in the two species relative to the radial axis. This is reminiscent of the inverted laminar organization of the cerebral cortex in mammals compared to other amniotes^21^.

The fraction of neurons classified as place cells varied strongly across recording locations, raising the possibility that place cells are anatomically organized within the hippocampus. We therefore systematically sampled throughout the volume of the hippocampus by varying the recording location across birds. We registered these locations to a template brain by constructing a three-dimensional model of the titmouse hippocampus (Extended Data Fig. 1). Both spatial information and spatial stability were strongly correlated to the recording location along the anterior-posterior axis (p < 10^-5^ for both measures; see Methods; **Fig. 2**), but not along the other two axes (p > 0.27 for both measures across both axes; Extended Data Fig. 7). Place cells were essentially restricted to the anterior two-thirds of the hippocampus, with incidence increasing from under 10% of excitatory cells at the posterior end to over 70% at the anterior end. This dramatic gradient is reminiscent of a similar specialization in the dorsal hippocampus of rodents^33^. In fact, the anterior-posterior axis of the avian hippocampus likely shares developmental origins with the rodent dorso-ventral axis (the “long” axis)^19–21,34^. Thus, not only the activity patterns, but even their anatomical organization, may be conserved across species.

Why did previous recordings in birds not reveal similar spatial representations^9–11,13^? One possibility is that place coding is species-specific, perhaps related to the ethological demands or experiences of a particular organism. As a result, place cells may be less common, less spatially precise, or more anatomically restricted in other species. To explore these possibilities while ruling out the effects of experimental technique, we repeated our random foraging recordings in the zebra finch — a species that, like those previously studied, is domesticated and does not cache food.

Systematic recordings across the zebra finch hippocampus revealed neurons with spatially modulated firing. As in titmice, place cells were found almost exclusively in the anterior hippocampus (Extended Data Fig. 8). Yet, in comparison to titmice, spatial activity was noticeably weaker and less abundant in the zebra finch. To quantify this difference, we sought to account both for a larger size of the titmouse hippocampus and for different sampling of neurons across the long axis in the two species. We therefore made comparisons in two ways, using landmarks defined either functionally or anatomically.

First, we defined an anterior segment of the hippocampus functionally, using the density of spatial cells in each species (see Methods). This segment was proportionately larger in titmice than in zebra finches (60% vs. 49% of the anterior-posterior extent of the hippocampus). Further, even within the anterior segment there was a greater density of place cells in titmice (60% vs. 47% of cells at the anterior pole; **Fig. 3a**). To illustrate this difference visually, we sorted cells in the anterior segment by spatial information to compare neurons with similar rank. More than 50% of these cells in titmice had spatial rate maps resembling those of classic place cells, whereas only about 15% did so in zebra finches (**Fig. 3b**).

Second, we identified a reliable anatomical landmark that divided the hippocampus roughly in half volumetrically (see Methods). We compared spatial information and spatial stability between species on the anterior and posterior sides of this divide. Both measures were significantly larger in titmice than in zebra finches in the anterior hippocampus (p = 0.002 and p = 0.0009, respectively), but not in the posterior hippocampus (p > 0.5; species difference was larger in anterior vs. posterior hippocampus, p < 0.01). Together, our analyses reveal a difference between species: place cells are more abundant and activity is more spatially precise and stable in titmice than in zebra finches. Such variability across species, as well as a dearth of place cells in a large posterior fraction of the hippocampus, might explain the reported absence of place cells in other non-food-caching birds^9–11,13^.

Our results so far show that “online” patterns of neural activity during active behavior are more similar between mammals and birds than previously thought. Might there be similarities in “offline” activity as well? In the mammalian hippocampus, periods of quiescence contain SWRs, defined by 1) a fast “ripple” oscillation in the local field potential (LFP), 2) a slower “sharp wave” deflection, and 3) synchronization of neural spikes to the ripple^4^. We examined activity during sleep in the avian hippocampus and indeed found events with these characteristics (in titmice: **Fig. 4a,b** and in zebra finches: Extended Data Fig. 8). SWRs were frequent (mean rate range 0.6−1.1 events/s in four titmice, n = 4−17 sessions, using conventional thresholds applied to the LFP in the 100−200 Hz frequency band; see Methods). As in rodents^31^, both excitatory and inhibitory cells increased firing during SWRs, but preferred different phases of the ripple oscillation (Extended Data Fig. 9). Thus, events resembling classic mammalian SWRs are indeed present in birds. In contrast, we did not observe continuous oscillations at any lower frequencies, including in the theta band, confirming that such oscillations are not universally required for spatial coding in the hippocampus (as in bats^35^; Extended Data Fig. 10).

A critical feature of SWRs in mammals is their global nature: events often extend across much of the hippocampal long axis, sometimes propagating in either direction^36^. We examined the organization of SWRs in titmice by implanting electrode arrays spanning >5 mm of the long axis. We found that individual events indeed often appeared to propagate across the hippocampus (**Fig. 4c**). About half of the events occurred on more than one electrode, with a sizeable fraction spanning most of the recorded extent of the hippocampus (length constant 0.90 mm; **Fig. 4d**). The median speed of propagation was 0.12 m/s (median absolute deviation 0.07 m/s, see Methods), and there was an abundance of events propagating in the posterior-to-anterior direction (70%, n = 15795 SWRs from 18 sessions from 2 birds; **Fig. 4d,e**), roughly consistent with the speed and directional bias observed in sleeping rats^36^. SWRs are therefore global events in the avian hippocampus, with organization across the long axis similar to that in mammals.

In mammals, the ability to detect SWRs is thought to depend on the crystalline cytoarchitecture of the hippocampus across the transverse plane (the plane perpendicular to the long axis)^37,38^. This includes a densely packed, planar pyramidal cell layer and parallel dendrites that allow small voltages to add together in the LFP. We were therefore surprised to observe SWRs in birds, whose hippocampus has only a modest clustering of cell bodies in a medially-restricted V-shaped region^26,27^ (**Fig. 1b**). Might there be hidden structure in the avian hippocampus that generates SWRs?

We noticed that as we lowered electrodes through the hippocampus, the sharp wave component often inverted from positive to negative polarity (**Fig. 4f**). To systematically examine SWRs across the transverse plane, we implanted additional titmice with arrays of electrodes oriented in the medial-lateral direction and recorded throughout the depth of the hippocampus (**Fig. 4g**). We then constructed two-dimensional current source density (CSD) maps to infer locations of sources and sinks (e.g. net positive current flowing out of or into cells, respectively)^37^. Remarkably, we observed a clear layered structure in the CSD map, with a current source located dorsal to a sink (**Fig. 4g,h**). Thus, there appears to be a laminar organization in the LFP of the titmouse hippocampus, in spite of dramatically different cytoarchitecture from mammals (**Fig. 4i**).

## Discussion

Place cells and SWRs are classic signatures of hippocampal function in mammals, but were unexpected in the avian brain. Birds are separated from mammals by 310 million years of evolution^30^, resulting in a hippocampus that is nearly unrecognizable at the levels of macroanatomy and cytoarchitecture^25^. Birds and mammals have divergent sensory systems, use these senses differently for navigation, and exhibit distinct methods of locomotion through the environment^39^. In fact, previous studies in non-mammals have not reported either hippocampal SWRs or precise spatial neurons akin to place cells^8–13^. Our results in such a distant system suggest that these activity patterns may be even more fundamental to memory functions of the hippocampus across amniotes (mammals, non-avian reptiles, and birds) than previously appreciated.

The anatomical organization of these activity patterns is equally surprising. We found two patterns of organization: a steep gradient of increasing place coding in the anterior direction, and a layered arrangement of current source densities during SWRs orthogonal to this direction. This organization is consistent with a conjectured homology of hippocampal axes across species: the anterior-posterior axis corresponds to the “long” dorso-ventral axis in rodents (posterior-anterior in primates), while the orthogonal direction corresponds to the radial axis^19–21,34^. In mammals these two functional axes are thought to result from organized anatomical features that are not apparent in birds: extrinsic inputs localized across the long axis^24^, and dramatic lamination across the radial axis^37,38^. Our results suggest that, in spite of these overt differences in anatomy, there are likely to be hidden, as-of-yet unidentified patterns of connectivity and cytoarchitecture that are conserved across species.

Place coding was present across two avian species, but was weaker in zebra finches than in titmice. There are many innate and experience-related differences between these birds (e.g. domestication), but it is tempting to speculate that precise spatial coding in titmice is related to the demands imposed by their food-caching behavior. Place cell activity is sparse^40^, and sparsity can allow neural circuits to form new memories quickly, while avoiding interference with old memories^40,41^. Since food-caching requires rapid memorization of new locations without forgetting old ones, increased sparsity of underlying neural circuits may confer an adaptive advantage to food-caching birds. However, the cost of sparse coding is a requirement for more neurons^42^. Thus, the benefit of sparsity in these species may have driven the enlargement of the hippocampus^28^, the prodigious capacity for spatial memory, as well as the abundance of spatially precise neurons that we found. Our results demonstrate a remarkable case of functional and anatomical conservation in a higher brain region of distant species. At the same time, these findings contribute to the growing body of evidence that hippocampal coding conforms to the unique ethological demands of different animals^43–49^.

## Acknowledgements

This work was supported by the Helen Hay Whitney Foundation Fellowship (H.L.P.), New York Stem Cell Foundation – Robertson Neuroscience Investigator Award, and the Beckman Young Investigator Award. We thank D. Scheck, S. Hale, T. Tabachnik, R. Hormigo, and K. Gutnichenko for technical assistance; M. Fee for the contribution to microdrive design; the Black Rock Forest Consortium, J. Scribner and Hickory Hill Farm, and T. Green for help with field work; L. Abbott and members of the Aronov lab for comments on the manuscript. The illustration of the arena in Fig. 1c and birds in Fig. 3a are by J. Kuhl.

## Extended data figures

**Extended Data Figure 1:**
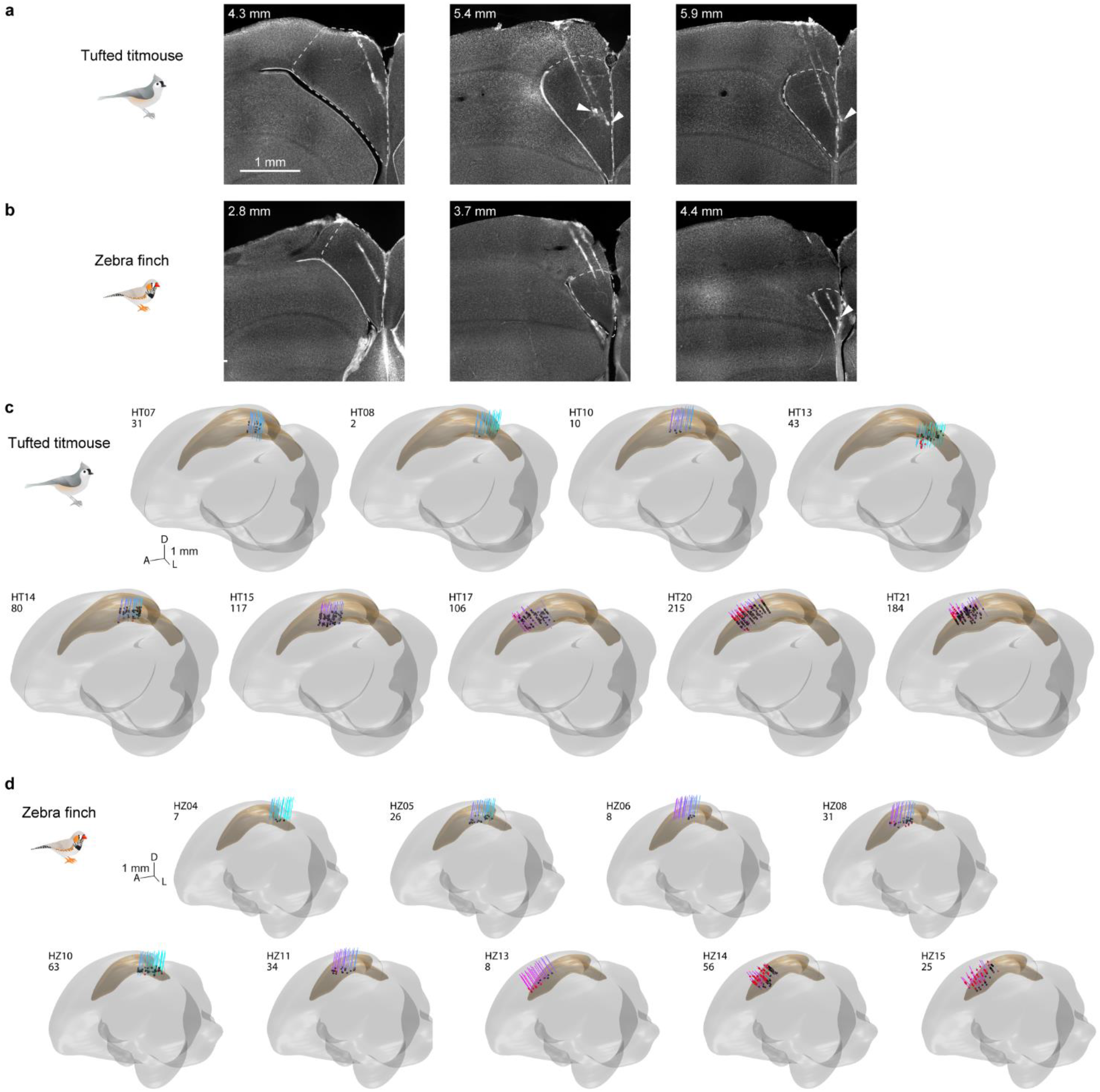
Histological mapping of recording locations. **a**, Fluorescent Nissl-stained coronal sections from a titmouse hippocampus. White arrowheads: lesion sites; dashed white: hippocampus outline. Numbers indicate anterior position relative to lambda. **b**, Same as in **a**, but for the zebra finch hippocampus. **c**, Reconstructed electrode tracks (blue lines, posterior; pink lines, anterior) for all nine titmice that were recorded during the random foraging task, aligned to a template brain. Grey: template brain; gold: template hippocampus. Black dots: locations of all cells recorded within the hippocampus; red dots: cells that were likely outside the hippocampus and were excluded from all analyses. Text indicates bird ID and number of hippocampal neurons recorded. **d**, Same as in **c**, but for zebra finches.

**Extended Data Figure 2:**
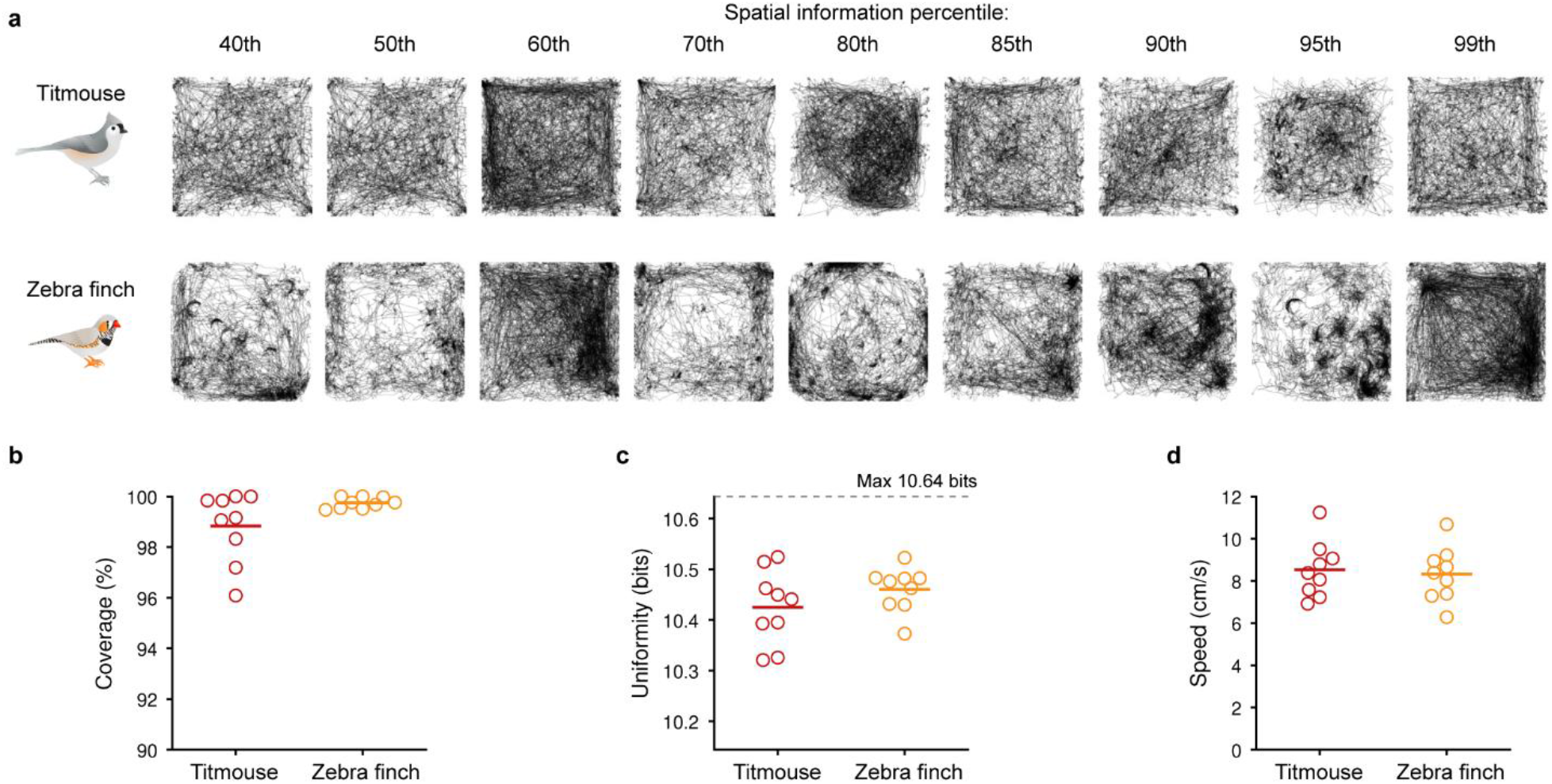
Both species achieve good arena coverage in the open field task. **a,** Behavioral trajectories for the sessions corresponding to the rate maps shown in **Fig. 3b**. **b**, Average coverage, measured as the fraction of bins in the smoothed spatial map bins that were occupied for at least 100 ms, and were thus included in the quantification of spatial information and spatial stability. Each marker represents the average coverage across all sessions for a single bird, in this panel and the following. Recording sessions with less than 90% coverage were not included in any analyses. **c**, Average uniformity of spatial occupancy maps, measured as − Σ*p_i_* log_2_(*p_i_*), where *p_i_* is the probability of occupancy in each spatial bin *i* (out of 40×40 = 1600 bins). The maximum possible uniformity is 10.64 bits (for a uniform distribution, where *p_i_* = 1/1600 for every *i*). **d**, Average speed, excluding stationary periods that were cropped from all analyses (see Methods).

**Extended Data Figure 3:**
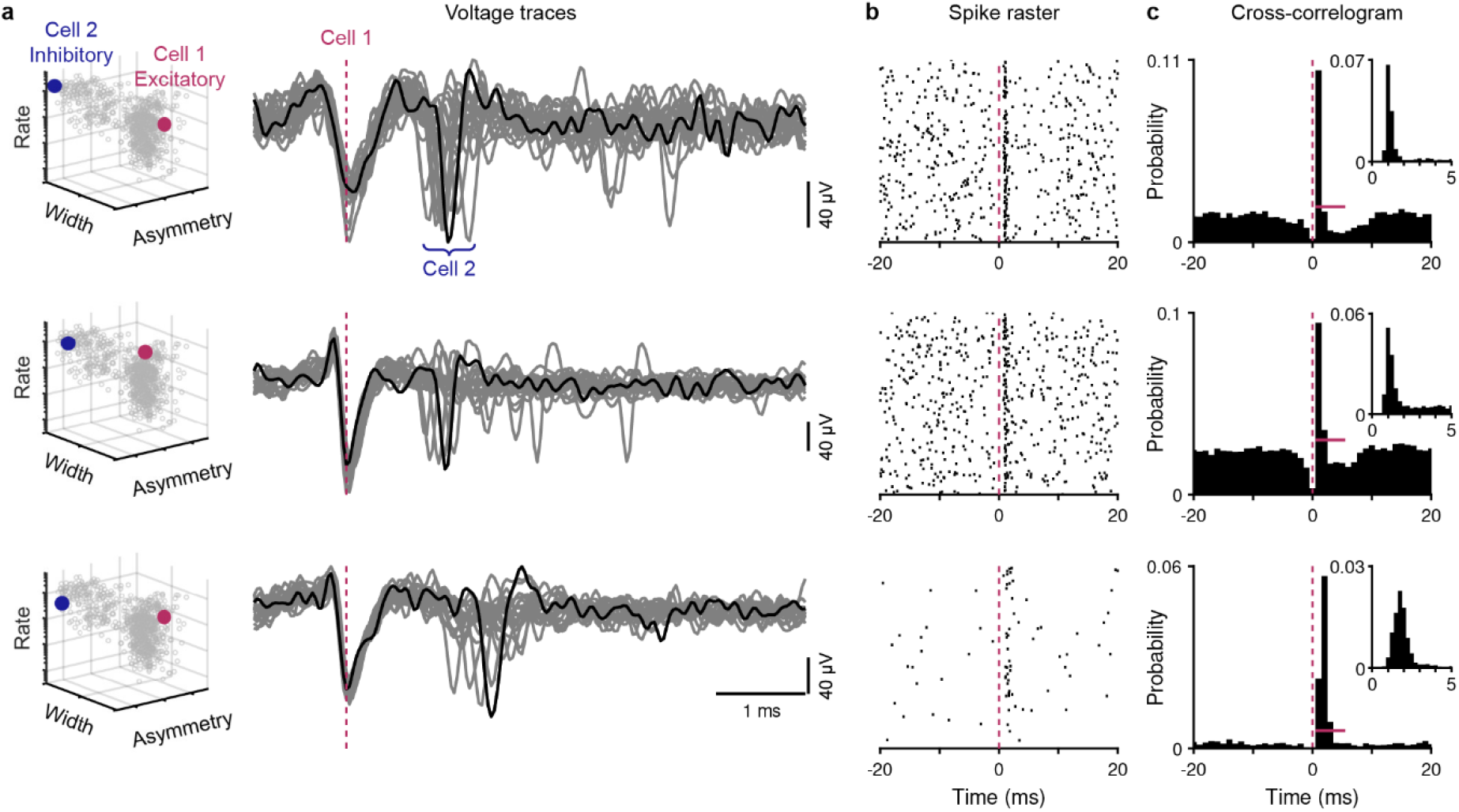
Short-latency spike correlations corroborate identity of putative excitatory and inhibitory neurons. **a**, Three pairs of putative excitatory and inhibitory neurons recorded on the same electrode. Left: each cell is highlighted in the plot of electrophysiological characteristics used to classify all cells as excitatory or inhibitory (replicated from **Fig. 1e**). Right: 20 consecutive voltage traces, spike-band filtered and aligned to the time of the excitatory cell spike. One example trace is highlighted in black. Note consistent short-latency responses in the inhibitory cell following excitatory cell spikes. **b**, Raster plot of the inhibitory cell spike times, aligned to 500 consecutive excitatory cell spikes (pink dashed line). **c**, Cross-correlogram between spike times in each pair of cells referenced to the excitatory cell (1 ms bins; inset: expanded histogram with 0.25 ms bins). The peak at positive latency indicates that inhibitory cell spikes tend to tightly follow excitatory cell spikes. The horizontal pink line designates the time span and threshold level used to assess significance of the putative monosynaptic excitatory interaction (see Methods). Of all 42 putative excitatory-inhibitory pairs, 24 pairs (57%) had significant excitatory interactions in the E→I direction, whereas none were significant in the I→E direction, in agreement with the putative identification based on waveform shape and spike rate (Fig. 1e). Additionally, as in mammalian cortex^1^, significant interactions were more likely for E→I pairs than for E→E pairs (5%, 2 of 38 pairs). The average latency for peaks in the cross correlation of significant E→I pairs was 1.7 ms (SD 0.7) (n = 23).

**Extended Data Figure 4:**
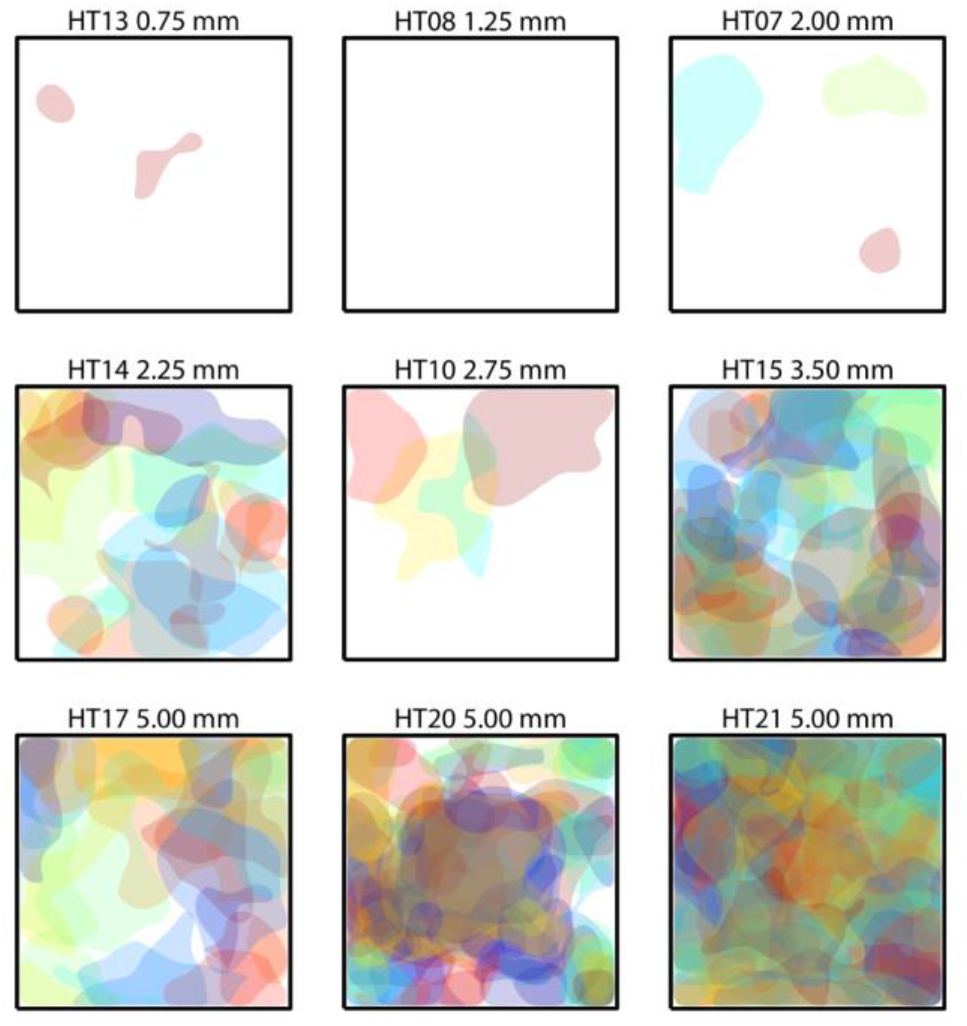
Firing fields of hippocampal neurons tile space. Significant firing fields for every place cell recorded in titmouse during the open field task. Each subpanel shows the fields recorded from a single bird. Bird ID and the average anterior position of the electrode bundle are indicated. Firing fields were defined as regions with areas of at least 72 cm^2^ (equivalent to a circle with diameter of 10 cm) where firing rate exceeded the 99th percentile of values for the corresponding bin from shuffled data. The boundary of each field was smoothed with a cubic smoothing spline (smoothing parameter 0.5). Note that some birds had few or no place cells either because few total cells were recorded and/or they were implanted at posterior locations. A random color was selected for each cell.

**Extended Data Figure 5:**
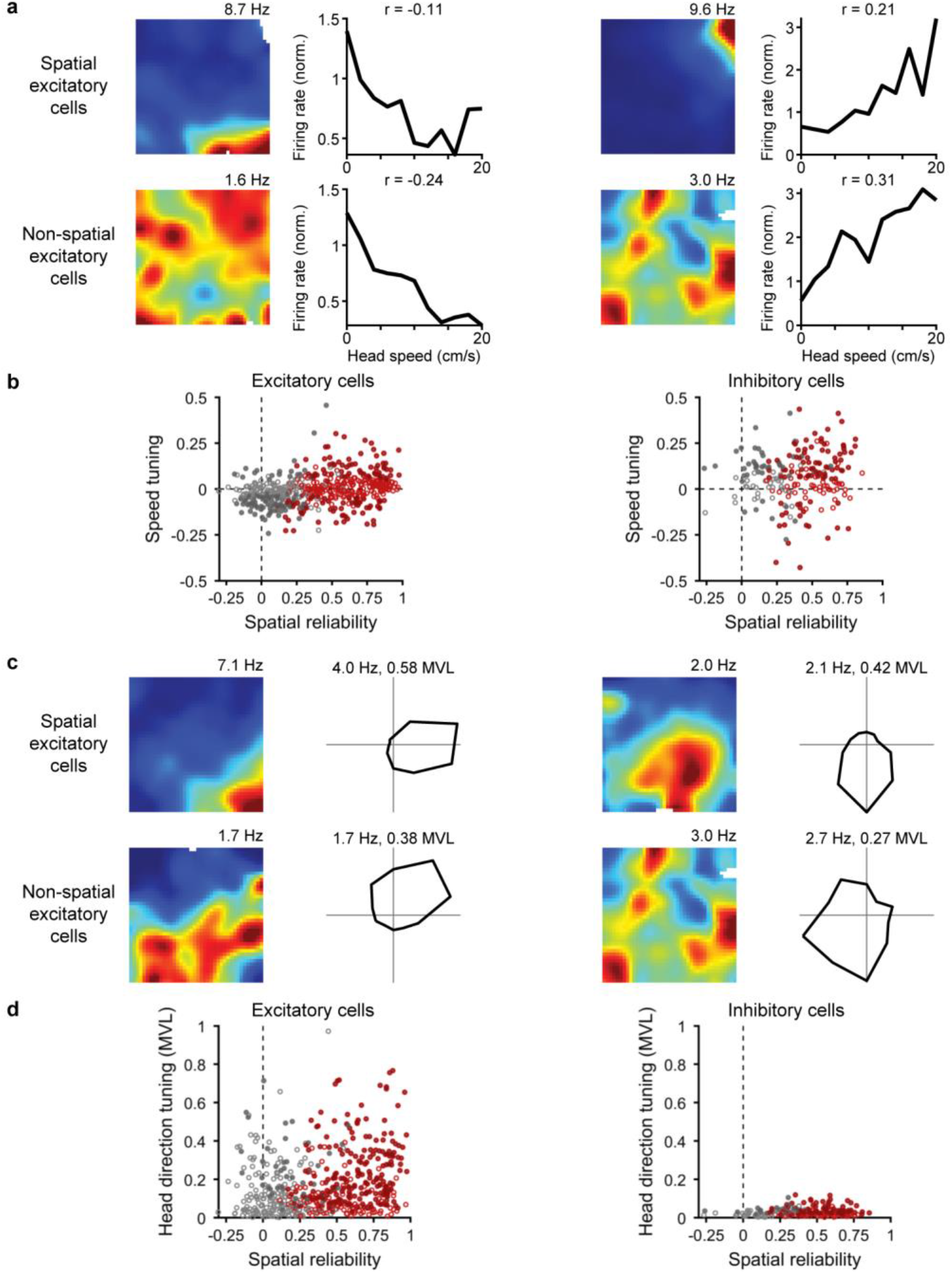
Tuning for speed and head direction in the titmouse hippocampus. **a**, Spatial rate maps and speed tuning curves for four example cells. Speed was binned for illustration purposes in these plots only. To calculate the speed tuning coefficient (r, noted above each plot), instantaneous speed was correlated with instantaneous firing rate. Top: place cells; bottom: non-place cells. Left: cells with negative correlation between speed and firing rate; right: cells with positive correlation. **b**, Population summary of speed tuning compared to spatial stability for excitatory and inhibitory cells. Red: place cells; grey: non-place cells; filled circles: significant speed tuning. Speed tuning was positively correlated with spatial tuning in excitatory cells; that is, cells that were less spatially reliable were more likely to have negative speed tuning, whereas cells that were more spatially reliable were more likely to have positive speed tuning (r = 0.28, p < 10^−10^). This correlation did not reach significance for inhibitory cells (r = 0.12, p = 0.09). Overall 15% of excitatory cells were positively correlated with speed and 27% were negatively correlated (n = 538). For inhibitory cells, 48% were positively correlated with speed and 12% were negatively correlated (n = 217). **c**, Spatial rate maps and head direction tuning plots for four example cells. Top: place cells; bottom: non-place cells. Maximum firing rate of the polar plot and the strength of head direction tuning measured by the mean vector length (MVL) are noted above (see Methods). **d**, Population summary of head direction tuning compared to spatial stability for excitatory and inhibitory cells. Colors as in **b**. MVL was correlated with spatial stability in excitatory cells (r = 0.21, p < 10^−5^); this relationship did not reach significance for inhibitory cells (r = 0.13, p = 0.052). Overall, 49% of excitatory cells (n = 522) and 45% of inhibitory cells (n = 210) had significant head direction tuning. Note that head angle was not available for a small number of recordings (16 excitatory and 7 inhibitory cells).

**Extended Data Figure 6:**
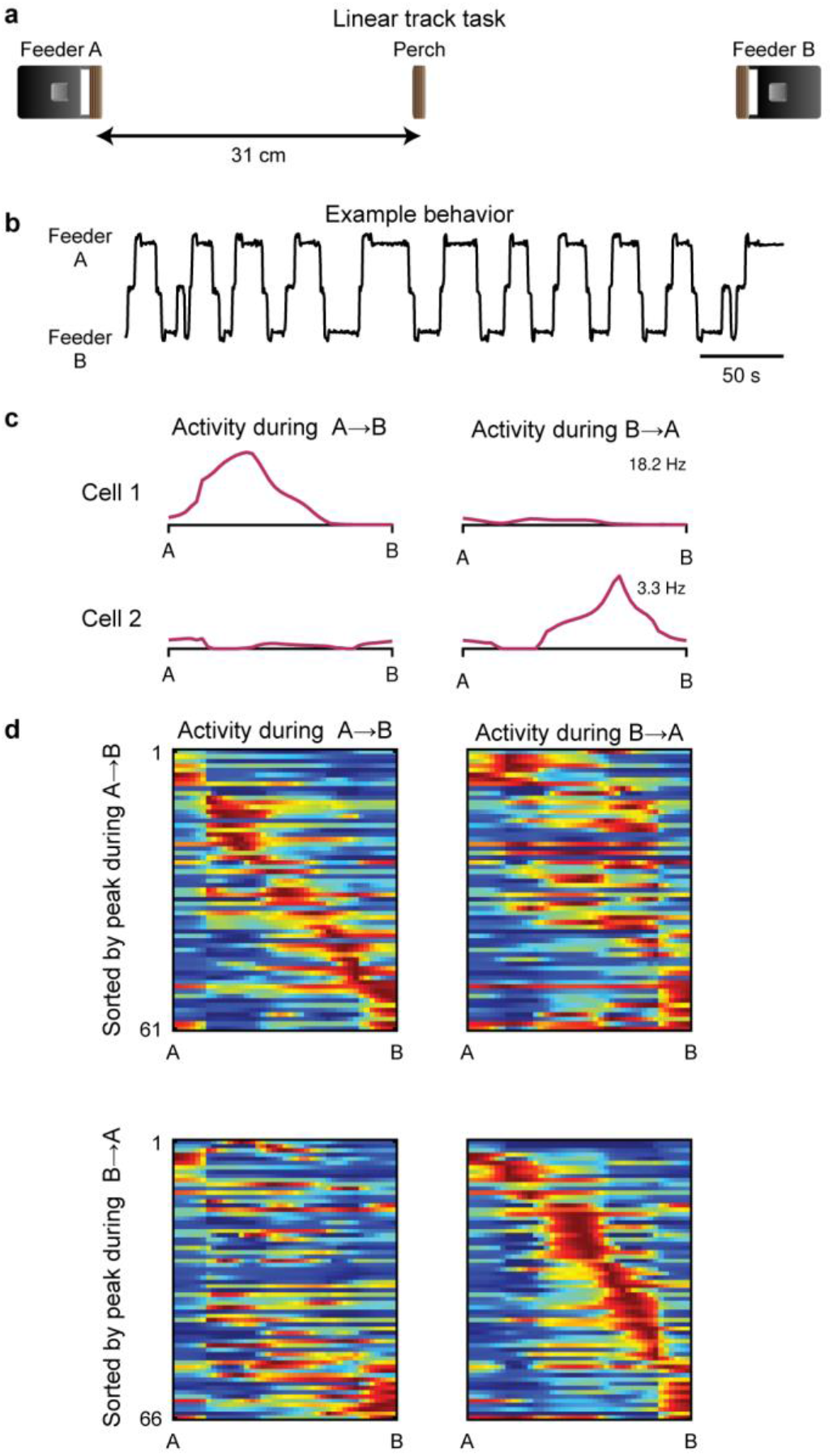
Directionally-selective place cells on a linear track. **a**, Schematic of task design. Three perches were arranged in a line, with motorized feeders at each end. Titmice learned to rapidly alternate between the two feeders over several days of training. **b**, Example behavioral trajectory. **c**, Firing rate as a function of position along the track for two example cells. One-dimensional rate maps were calculated separately for each direction of travel. The peak firing rate for each cell is indicated. Note direction selectivity. **d**, One-dimensional rate maps for all excitatory cells recorded on the linear track. Each row represents a single cell. Left and right columns show rate maps calculated for movement during each of the two directions. Color scale: blue, 0 Hz; red: 99th percentile of firing rate. Top: only cells that had significant firing fields in the A→B direction (n = 61 of 105), sorted by the location of the peak in the firing rate map for that direction. Bottom: only cells that had significant firing fields in the B→A direction (n = 66 of 105), sorted by the location of the peak in the firing rate map for that direction. Of the 105 cells, 49 had significant firing fields in both directions.

**Extended Data Figure 7:**
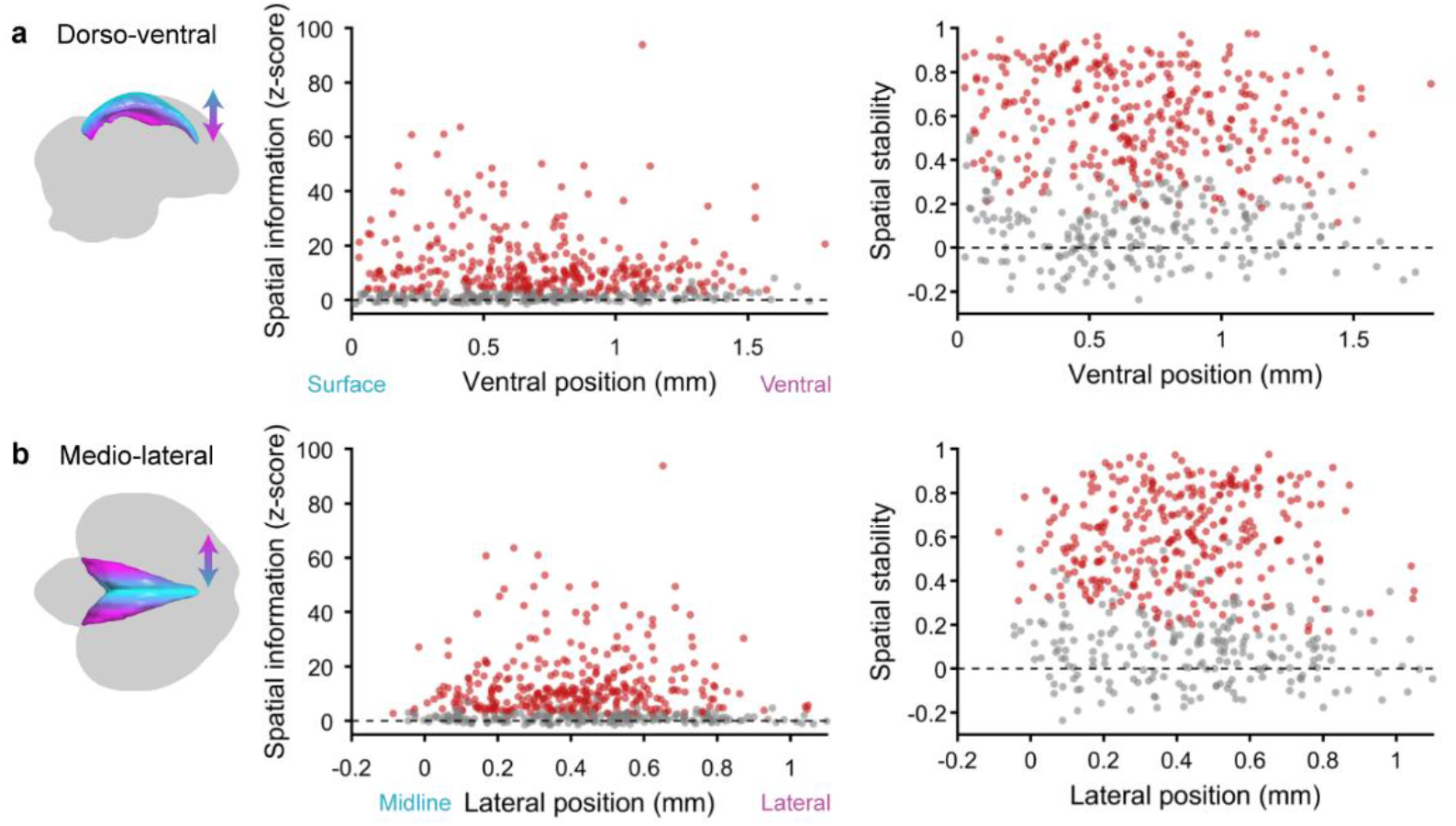
Lack of spatial tuning gradients across the dorso-ventral and medio-lateral axes of the titmouse hippocampus. **a**, Spatial information and spatial stability plotted against ventral position, measured relative to the dorsal-most position of the hippocampus in each coronal section. Colors as in **Fig. 2 b,c**. **b**, Spatial information and spatial stability plotted against the lateral position of the recording, measured relative to the midline. Some cells have negative medio-lateral positions due to variations of hippocampal shape relative to the template brain.

**Extended Data Figure 8:**
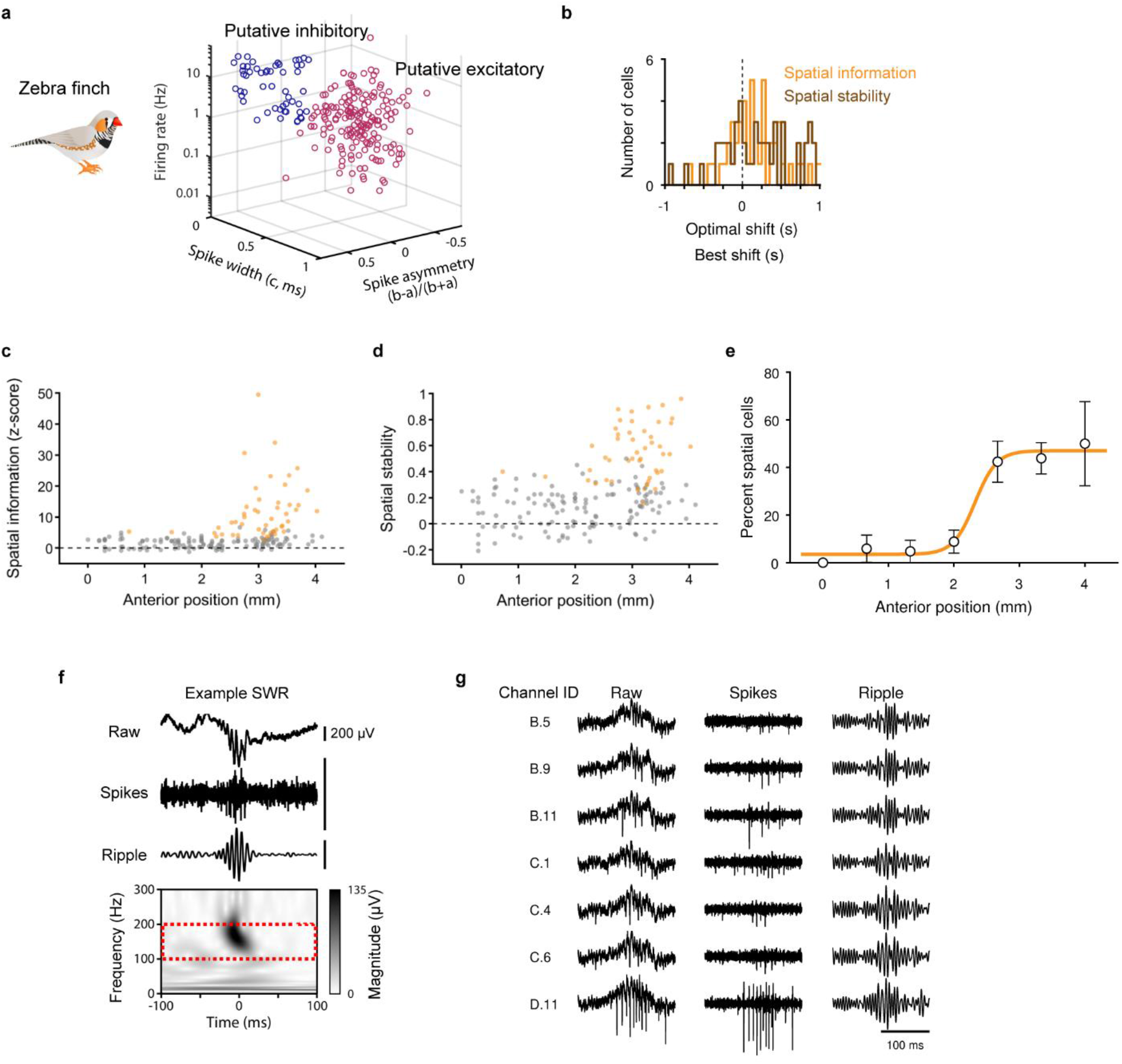
Spatial tuning and SWRs in the zebra finch hippocampus. Results in the zebra finch, similar to those shown in the main figures for the titmouse. **a**, Putative excitatory (n = 179) and inhibitory cells (n = 59) during the open field task, as in **Fig. 1e**. **b**, Optimal time shifts for spatial information and spatial stability, as in **Fig. 1g**, bottom (mean optimal shift 144 ms and 166 ms, respectively; both greater than zero, p = 0.005 and 0.012, two-sided t-test, n = 47 and n = 48). **c**,**d** Normalized spatial information and spatial stability increased with anterior position (p = 0.013 and p = 0.0007, respectively, likelihood ratio test of linear mixed effects model), as in **Fig. 2b,c**. **e,** Fraction of place cells as a function of anterior position, as in **Fig. 2d**. **f**, Example SWR, as in **Fig. 4a**. **g**, Example SWR recorded across seven contacts of a silicon probe in the zebra finch hippocampus. Recordings are from three probe shanks, with inter-shank spacing of 200 μm. Shank (B, C, D) and contact number are noted.

**Extended Data Figure 9:**
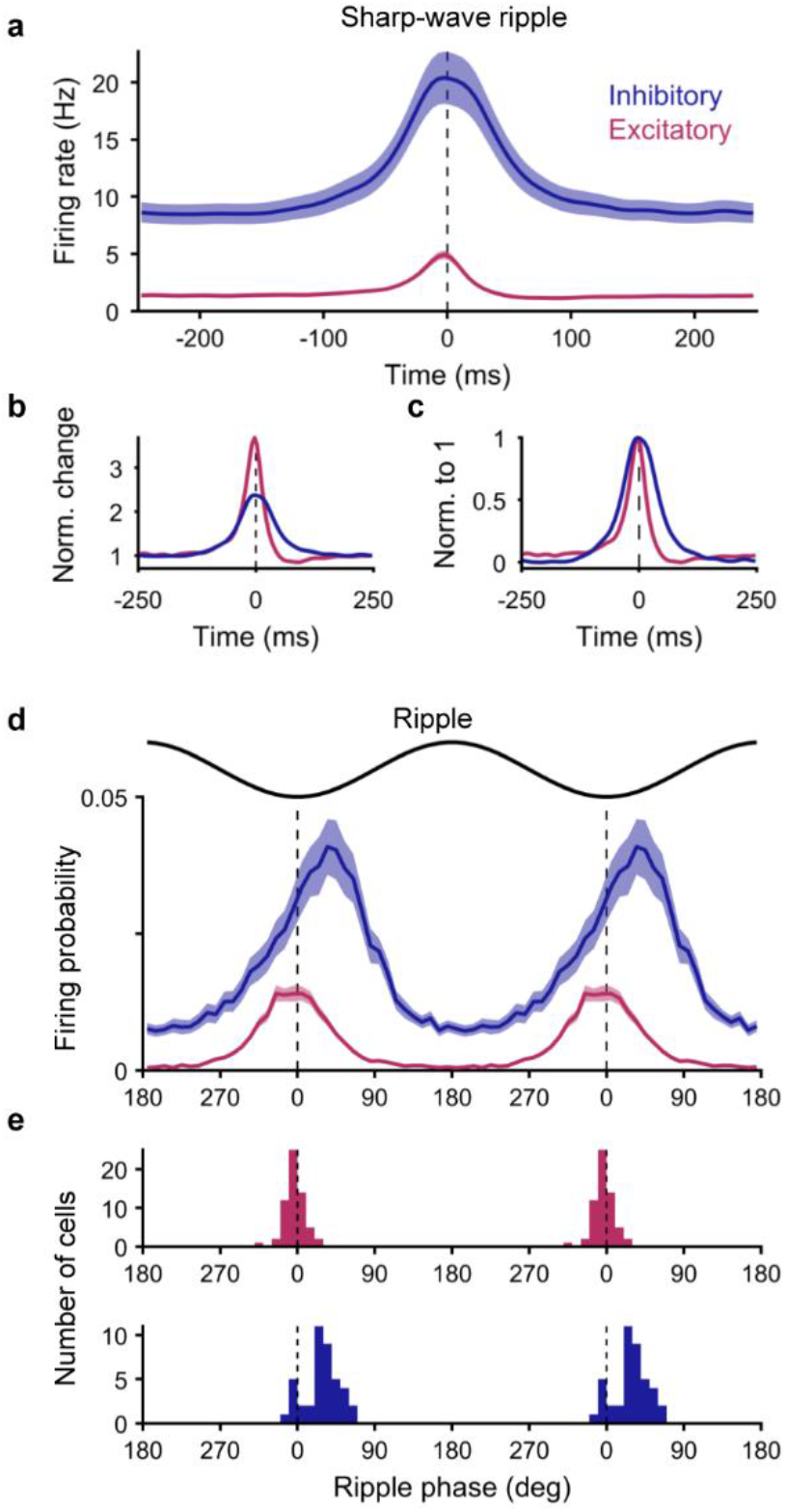
Modulation of spiking in excitatory and inhibitory neurons during SWRs. **a**, Average firing rate of excitatory (n = 65) and inhibitory (n = 43) neurons during SWRs in sleeping titmice. Both excitatory and inhibitory neurons increased their firing during SWRs, as in rodents^2^. Error bars: mean ± SEM. **b**, Same as in **a**, but normalized by the average baseline firing rate from ±150–250 ms. As in rodents^2^, excitatory cells showed a larger proportional increase in firing rate than inhibitory cells (3.7-and 2.4-fold, respectively). **c**, Same as in **a**, but normalized from 0 to 1. As in rodents^2^, inhibitory cells showed broader temporal activation. This could potentially arise from a combination of unidentified subclasses of inhibitory cells. **d**, Average firing probability of all excitatory and inhibitory neurons across phases of the ripple cycle. Only spikes occurring during the SWR, defined as the period over which the ripple envelope exceeded two standard deviations of baseline, were included. For visualization only, the result was duplicated and concatenated across two ripple cycles. Error bars: mean ± SEM. **e**, Histogram of preferred ripple phase for excitatory (top) and inhibitory (bottom) cells. Preferred phase was determined by fitting a sine wave to the ripple-aligned spike probability histogram for each cell. Only cells with significant fits were included in this panel. Excitatory cells reliably fired near the trough of the ripple (mean −3.1°), whereas inhibitory cells tended to fire after the trough (27.6; difference between excitatory and inhibitory p < 10^−15^), similar to rodents^2^.

**Extended Data Figure 10:**
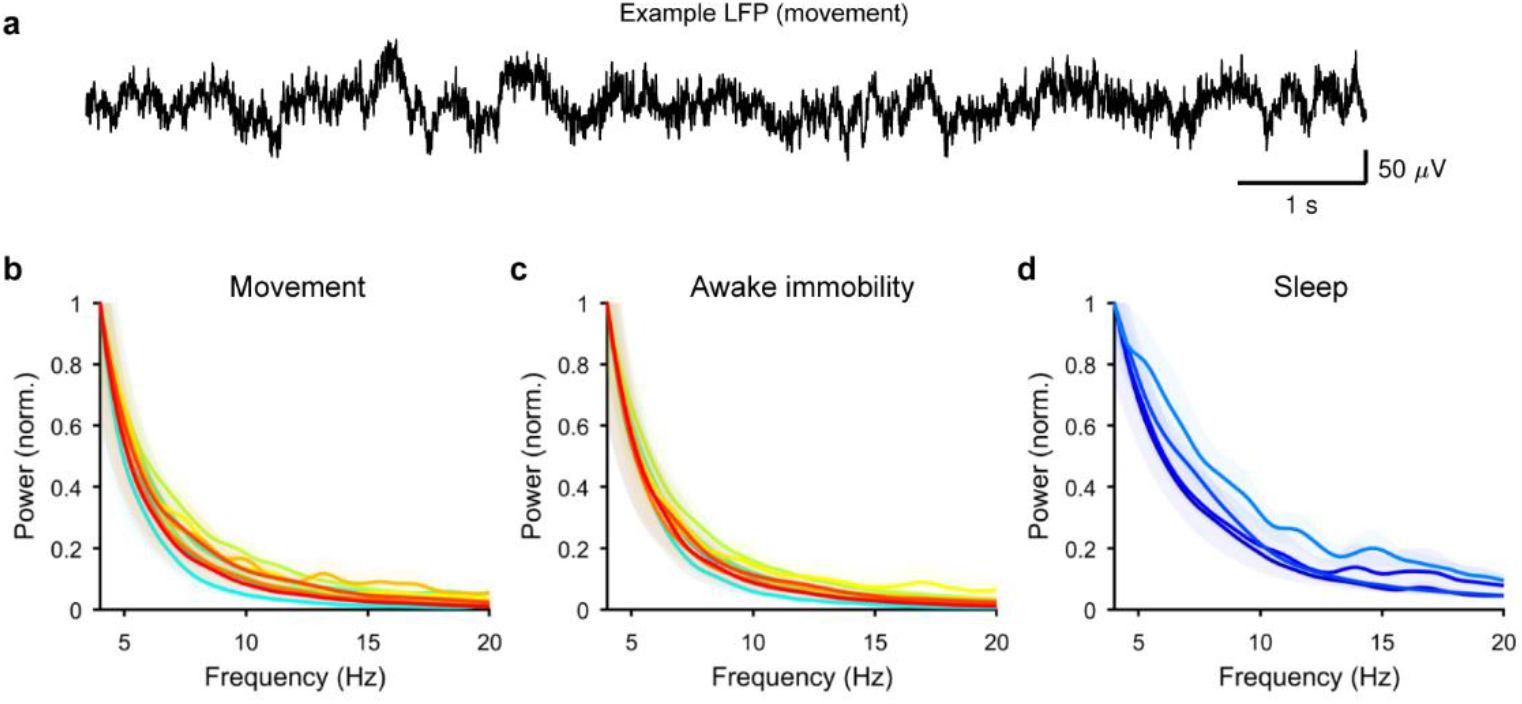
Lack of evidence for theta oscillations in the bird hippocampus. **a**, Typical example of raw LFP during the open field task, low-pass filtered at 100 Hz to remove spikes. **b**, Power spectra of the LFP during movement in the open field task. Each line represents the average power spectrum for a single bird (n = 9 titmice, error bars: mean ± SEM across sessions). **c**, Periods of immobility in the open field task, plotted as in **b**. **d**, Recordings during sleep, plotted as in **b** (n = 4 titmice).

## Methods

### Subjects

All animal procedures were approved by the Columbia University Institutional Animal Care and Use Committee and carried out in accordance with the US National Institutes of Health guidelines. Subjects were 15 adult tufted titmice (*Baeolophus bicolor*, collected from multiple sites in New York State using Federal and State permits) and 11 adult zebra finches (*Taeniopygia guttata*, Magnolia Farms). The open field experiments include data from 9 tufted titmice (5 female, 4 male) and 9 zebra finch (all male). The linear track experiment includes data from 3 tufted titmice (all female). SWRs were analyzed during sleep in 4 tufted titmice (2 female, 2 male) and 2 zebra finches (1 female, 1 male). Finally, one subject of each species was used to construct a 3D model of the brain and hippocampus. There is no visible sexual dimorphism in tufted titmice, so experiments were blind to sex, which was determined after perfusion. Only male zebra finches were used because they are generally larger, and more easily support the weight of the microdrive and cable during open field behavior. All birds were singly housed and subject to a “winter” light cycle, 9h:15h light:dark. Random foraging and linear track experiments were run during the light ON period, and SWR recordings during sleep were run during the light OFF period. Primary wing feathers were trimmed to prevent flight.

### Behavioral experiments

Experiments in awake birds were conducted in an enclosed square arena, with a central open space 61 cm on each side, partially surrounded by a 2.5 cm boundary interrupted by corner posts. The walls, floor, and ceiling were black, with ~15 cm diameter bright shapes (yellow circle, pink star, blue pentagon, and green tree) positioned on each wall, centered ~30 cm above the floor. The arena was illuminated from above. White noise was played in the background to mask inadvertent room noises. Position was tracked using an infrared motion-tracking system (Qualisys) at 300 frames/s. Sessions typically lasted 1 hour, and each bird was run once per day.

To motivate consumption of seeds during the open field and linear track tasks, birds were deprived of food from the time of light onset until the start of the experiment. Birds were weighed daily before the experiment. The time period of food deprivation after light onset ranged from 1–5 hours, adjusted to achieve stable weight.

In the random foraging task, small sunflower seed fragments (2.5 mg, for titmice) or millet seeds (1.3 mg, for zebra finch) were automatically released from above by a custom-built automatic seed dispenser. These food items were dropped from a sufficient height (110 cm) that they scattered randomly and virtually uniformly after hitting the floor. The rate of the seed dispenser was adjusted to roughly match the pace at which the birds consumed seeds. Birds typically underwent 3–6 habituation sessions, some conducted before surgery and some afterwards. Electrophysiological recordings began after these sessions.

In the linear track task, two custom-built feeders were positioned at opposite corners of the arena. These feeders were equipped with motion sensors to detect when the bird retrieved a seed. When a seed was retrieved from one feeder, that feeder closed and the other feeder opened. Birds were motivated to alternate between the feeders, and reliably learned to do so after several days of pre-training.

For recordings of SWRs during sleep, birds were recorded in the dark in a smaller chamber with a central perch, placed inside the larger arena. The bird was monitored with an infrared video camera. It was considered asleep when it exhibited characteristic fluffed feathers, head tucked under a wing, and lack of motion. Several recording sessions of 1 hour each were conducted consecutively each night, with incremental advancement of the electrodes before each session.

### Electrophysiology

Neural recordings were obtained using custom miniature microdrives. Each microdrive contained an array of platinum-iridium microelectrodes (Microprobes PI20035.0A3 and PI20031.0A3). Twenty-one electrodes with 5 MOhm impedance were used for random foraging and linear track experiments, and 7–13 electrodes with 1 MOhm impedance were used for recordings during sleep. (For recordings during sleep in one zebra finch (Extended Data Fig. 8g), a silicon probe (Cambridge NeuroTech, ASSY-236F) was used instead.) Each microelectrode was guided through polyimide tubes (ID 0.004”, OD 0.055”; Nordson medical #141-0001) arranged in a bundle. The electrodes were then soldered to a flexible cable attached to a moveable shuttle, allowing the bundle of electrodes to be advanced into the brain by turning a single screw (size M1.2-0.25 x 10mm). Typically, the drive was advanced immediately before each recording session by 1/8th or 1/4th of a full 250 μm turn. A protective housing was built using 3D printed polymer and transparency film and shielded using conductive paint (Ted Pella, 16062). The dimensions of the microdrive were 12 mm wide × 6 mm deep × 24 mm high, and the total weight including cement was typically 1.8 g for titmice and 1.6 g for zebra finches. Passive IR-reflective markers for motion tracking added 0.6 g for titmice and 0.2 g for zebra finches.

Signals were referenced to a platinum-iridium wire (0.002” diameter bare, 0.004” coated, AM Systems 776000) implanted outside of the hippocampus in the same hemisphere. A silver ground wire (0.005” diameter bare, AM Systems 781500) was implanted in the opposite hemisphere.

Signals were amplified, multiplexed, and digitized at 30,000 samples/s within the microdrive housing using an interface chip (Intan Technologies, LLC; RHD2164) wire-bonded onto a custom PCB. Digital signals were then transmitted from the bird to a computer interface board via a 12-conductor SPI cable (Intan technologies, LLC; C3213), with outer sheathing stripped to reduce weight. The effective weight of the cable was further reduced by attaching a thin piece of very elastic material to either end (Linsoir Beads, Crystal String). A motorized commutator (Doric Lenses, Inc., AERJ_24_HDMI) allowed the bird to turn freely.

### Surgery

Birds were anesthetized using 1.5% isoflurane in oxygen and injected intraperitoneally with dexamethasone (2 mg/kg). Feathers were removed from the surgical site, and birds were placed in a stereotaxic apparatus using ear bars and a beak clamp. To anchor the microdrive, 4–6 partial craniotomies through the outer layer of the skull were filled with light-cured dental cement (Pentron Clinical, N11H). A full craniotomy and durotomy were made above the hippocampus at the recording locations noted in the text. For two birds in which SWRs were recorded across the transverse plane, electrodes were implanted 3 mm anterior to lambda. All microdrives were implanted into the left hippocampus, tilted to the left by 20° in order to target the medial hippocampus while avoiding the large blood vessel over the midline. For most birds, the brain was tilted such that the angle of the groove at the base of the upper mandible of the beak was 65° relative to the horizontal. For the anterior-most recording locations (titmice HT17, HT20, HT21 and zebra finch HZ13, HZ14, and HZ15 in Extended Data Fig. 1c,d), electrodes were implanted with the brain tilted up by an additional 30° to allow more direct access to the hippocampus, which is located below the surface of the brain in this anterior region. Buprenorphine (0.05 mg/kg) was injected intraperitoneally after the surgery.

### Histology

After completion of the experiment, electrode locations were marked by passing 30 μA negative current for either 1 s or 5 s through each electrode. Animals were given an overdose of ketamine and xylazine and were then perfused transcardially with saline followed by 4% formaldehyde. Brains were extracted and stored in 4% formaldehyde, then cut into 100 μm-thick coronal sections. Brain sections were either stained with fluorescent Nissl stain (Invitrogen N21480), fluorescent DAPI (Invitrogen D1306), or left unstained. For the three titmice and three zebra finches that had microdrives implanted in very anterior hippocampus, where the hippocampus is submerged below the dorsal surface of the brain (birds HT17, HT20, HT21, HZ13, HZ14, and HZ15 in Extended Data Fig. 1c,d), as well as for the two titmice in which SWRs were recorded across the transverse axis, the position of each electrode relative to the boundary of the hippocampus was reconstructed using lesion sites and electrode tracks. This reconstruction was used to determine which recorded cells were likely outside the hippocampus; these cells were thereby excluded. For other birds, the stereotaxic coordinates of the implant site were registered to a template brain, and the small number of cells that were estimated to be outside the hippocampus were excluded (Extended Data Fig. 1c,d).

To facilitate both surgical planning and registration of final recording locations, we constructed template brains containing a 3D reconstruction of the hippocampus aligned to stereotaxic coordinates. For each species, stiff wire (Malin Co., 0.006” music wire) was inserted into the brain horizontally at two locations during surgery. Stereotaxic coordinates of both locations were recorded. The brain was then sectioned sagittally. The outline of the hippocampus and the whole brain was traced in each section using Matlab and Illustrator. Visible holes made by the wire allowed registration of sections to each other and to stereotaxic coordinates.

Compared to neighboring regions, the hippocampus in birds has a lower density of cells and larger cell bodies, making the identification of hippocampal boundaries unambiguous^3^. This definition of the hippocampus in birds appears to be homologous to the hippocampus proper in mammals, i.e. Ammon’s horn and the dentate gyrus, based on genetic analysis^4–6^.

### Behavioral analysis

Titmouse behavior was tracked using a rigid arrangement of infrared-reflective markers connected to the microdrive, which allowed 6DOF tracking of head position by the software QTM (Qualisys). For the analyses in Figs. 1 and 2, the location of the titmouse was defined as the midpoint between the eyes projected onto the floor of the arena. The location of the midpoint between the eyes was determined by measuring the positions of the markers relative to the eyes of the bird, either during surgery or after perfusion. For zebra finches, only a single marker was affixed at the top of the microdrive in order to reduce the weight of the implant. The position of this marker projected onto the floor was used as the position of zebra finch in the open field arena. For direct comparisons between titmice and zebra finch (Fig. 3), an equivalent position relative to the microdrive was used as the position of the titmouse in the arena.

Movement speed was calculated by first smoothing x and y position separately with a 1-s moving average, then differentiating each and calculating absolute speed as 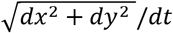. This smoothing window was chosen to reduce the influence of head movements, and to instead capture locomotion through the arena. We excluded stationary periods when the speed fell below 5 cm/s for longer than 5 s from all analyses of spatial tuning and head direction tuning.

### Spike sorting

Single units were identified using Plexon spike sorting software. Raw signals from individual electrodes were first high-pass filtered using a 4-pole Butterworth filter at 250 Hz. Candidate waveforms were thresholded, projected into 2D or 3D PCA space, and then sorted into putative single units using template sorting. For display purposes, spike waveforms and voltage traces shown in the figures were band-pass filtered from 800–5000 Hz (Hamming window FIR filter).

### Identification of excitatory and inhibitory neurons

All analyses were conducted in MATLAB. Putative excitatory and inhibitory neurons were identified by applying k-means clustering with two clusters to the electrophysiological characteristics described in Fig. 1e. Spike width was calculated as the time from the trough of the average spike waveform to the subsequent peak. Peak amplitude asymmetry was calculated as the relative height of the two positive peaks flanking the trough (Fig. 1e)^7^. This measure equals −1 when the first peak is present and the second is non-existent, 0 when the two peaks are equal, and +1 when the second peak is present and the first is non-existent. Mean firing rate was 1.2 Hz and 11.5 Hz in the putative excitatory and inhibitory clusters, respectively; spike width was 0.60 μs and 0.31 μs; and peak amplitude asymmetry was −0.21 and +0.47.

We further assessed our criteria for classifying excitatory and inhibitory neurons by examining spike trains of pairs of neurons recorded on the same electrode. Putative monosynaptic connections between neurons have been inferred using cross-correlograms of spike times^1,8^. For each pair of cells, we constructed the cross-correlation between spike trains binned to 1 ms resolution with time lags of ±20 ms. Only cells with at least 1000 spikes were included. To test for significance, we also constructed jittered cross-correlograms by shifting each spike time by a random value drawn from a uniform distribution in the interval ±5 ms. This interval was intended to preserve slower, non-monosynaptic spike time correlations, such as those arising from common input. Our spike sorting method precluded detection of spikes from two different neurons recorded on the same electrode within ~1 ms of each other, so any spikes from the two neurons that fell within the same 1 ms bin were re-jittered until there were no such overlaps. The maximum value of the jittered cross-correlogram within the interval from 1 ms to 4 ms was stored, and this process was repeated 500 times to create a null distribution. A putative excitatory monosynaptic connection was considered significant if there was a peak in the actual cross-correlogram within the 1-4 ms interval, and if this peak ms exceeded 99% of the null-distribution values. We only assessed putative excitatory connections because the cross-correlogram technique is much more sensitive to excitatory interactions than inhibitory ones^9^.

Unless otherwise specified in the text (e.g. Fig. 1 d-f, Extended Data Fig. 5, Extended Data Fig. 9), all analyses were conducted on excitatory cells only.

### Analysis of spatial activity in the random foraging task

Spatial firing rate maps were constructed to capture the relationship between neural activity and spatial location. The arena was divided into a grid of 40×40 bins. For each neuron, spike counts and occupancy time in each bin were calculated, and the resulting matrices were separately smoothed with a 13×13-point Hamming window. Any bin with less than 0.1 s occupancy after smoothing was replaced with “Not a number” (NaN). Any cell with poor coverage, defined as more than 10% of the arena occupied by NaN, was excluded from further analysis. We also excluded cells for which the total distance travelled during the recording was less than 50 m. Finally, spike counts were divided by occupancy to yield mean firing rates in each bin.

We quantified the degree of spatial tuning for each neuron in two ways. First, we calculated the amount of information about position conveyed by firing rate according to:

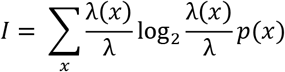

where *I* is the information rate in bits/spike, *x* is the spatial bin, *p*(*x*) is the probability that the bird occupies bin *x*, λ(*x*) is the mean firing rate in bin *x*, and λ = Σλ(*x*)*p*(*x*) is the overall mean firing rate^10^. Higher information rates arise from sparser spatial rate maps, indicating that each spike is highly informative about position. Because spuriously high spatial information values can arise from cells that only fire a small number of spikes, spatial information was normalized by taking the z-score relative to the distribution for a shuffled dataset (see below).

Second, we calculated the stability of the spatial rate map over time^11,12^. The recording session was divided into 5-min, non-overlapping segments. Half of the segments were randomly assigned to set 1, and the other half to set 2. Spatial rate maps were constructed separately for each set, and the correlation between the two maps was calculated. This process was repeated ten times, and the results were averaged to yield a final correlation value. Higher correlations indicate greater stability of the spatial rate map.

To assess significance of spatial coding for each cell, both of the above measures were also calculated for shuffled data. For each iteration, spikes were circularly shifted relative to behavior by a random time delay from 0 to the duration of the recording session. Spatial information and spatial stability values were calculated and stored. This process was repeated 200 times to construct a distribution of values representing the null hypothesis for each cell. Only cells for which both spatial information and spatial stability exceeded the 99th percentile of the corresponding shuffled distribution were considered significant.

We calculated spatial information and spatial stability at different time shifts between spikes and behavior, from –1 s to +1 s. Only spatial excitatory cells that had peak values within this time range were included in the statistical tests.

To assess the significance of spatial coding across the long axis of the hippocampus, we used a linear mixed effects model, which is well-suited for datasets with multiple observations per subject — here, multiple cells recorded from each bird (Oberg & Maloney 2007). We used the following model to test whether anterior position predicted spatial stability and/or normalized spatial information:

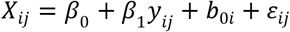

where

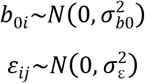

and *X*_*ij*_ is the variable of interest (spatial stability or information) for the *j*^th^ cell from the *i*th bird, *β*_0_ is the fixed intercept, *β*_1_ is the fixed effect coefficient for anterior position, *y_ij_* is anterior position, *b*_0*i*_ is the random intercept for each bird, and *∊_ij_* is the residual error. Because normalized spatial information across cells was not normally distributed, we applied a Yeo-Johnson transformation with λ = 0 to reduce positive skewness before applying the linear mixed effects model^13^. The model was then fit using the *fitlme* function in MATLAB using the default maximum likelihood method. Significance was determined by calculating the log-likelihood of the full model compared to a reduced model with the fixed slope *β*_1_ removed using the *compare* function.

An extended model was constructed to assess spatial tuning across dorso-ventral and medio-lateral axes (Extended Data Fig. 7):

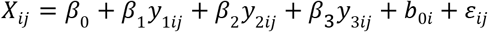

where *β*_1_, *β*_2_, and *β*_1_ are the fixed effect coefficients for anterior, lateral, and ventral position, respectively, y_1ij_, y_2ij_, and y_3ij_ are the corresponding positions, and other variables are as above. Significance was assessed by removing one fixed effect at a time, and comparing the full model to this reduced model by taking the log-likelihood. Anterior position was significant in this extended model (p = 0.001 and p = 10^−5^ for spatial information and spatial stability, respectively).

### Analysis of speed and head direction tuning

Speed tuning was assessed by calculating the correlation coefficient between speed and firing rate. Speed was calculated as described above, and firing rate was calculated in a 1-s moving window. The resulting correlation coefficient was deemed significant if it exceeded 99.5%, or was below 0.05%, of the distribution for the shuffled dataset, in which spikes were randomly shifted relative to behavior (see above).

Head direction was calculated as the component of head angle (the vector from the point between the eyes to the tip of the beak) projected onto the plane of the arena floor. We binned head direction into 36° bins and calculated the average firing rate in each bin. The resulting values were used to compute the mean vector length (MVL) for the polar rate map of head direction^11^. MVL quantifies the strength of head direction tuning. A MVL equal to 0 indicates that firing rate is uniform across heading directions, whereas a MVL equal to 1 indicates perfect tuning to a single bin of head direction, and no firing for other directions. Head direction tuning was considered significant if the real MVL was larger than 99% of the MVL for the shuffled dataset.

### Comparison of spatial activity in zebra finches and titmice

To compare neural characteristics of anterior and posterior hippocampus across avian species, we defined both functional and anatomical landmarks separating anterior and posterior hippocampus. Note that we do not imply any sharp division between anterior and posterior hippocampus. Instead, the goal was to allow comparison between similar regions across species. The anatomical landmark may be additionally useful for future analyses of other avian species, for which systematic recordings across the long axis are not available.

The functional landmark was defined as the inflection point of a logistic sigmoid function fitted to the percentage of spatial cells binned according to anterior position. The bins were 1 mm for titmice and 0.67 mm for zebra finch, resulting in seven bins for each species. The region anterior to the functional landmark thus displayed a relatively high density of spatial cells in both species.

To define the anatomical landmark, we selected all transverse sections of the brain in which the hippocampus contacted both the dorsal brain surface and the midline. Anterior to these sections, the hippocampus was ventral to the hyperpallium and did not contact the brain surface. Posterior to these sections, the hippocampus was lateral to the cerebellum and did not contact the midline. The anatomical landmark separating anterior and posterior hippocampus was the location halfway between the anterior-most and the posterior-most selected brain sections. We found that roughly half of the hippocampal volume was anterior to this location, and half was posterior.

We used a linear mixed effects model to assess the significance of differences in spatial coding between species (similar to that described in the section above “*Analysis of spatial activity in the random foraging task”*). The model was as follows:

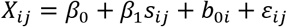

where *X_ij_* is the variable of interest (spatial stability or normalized information with Yeo-Johnson transformation) for the *j*th cell from the *i*th bird, *β*_0_ is the fixed intercept, *β*_1_ is the fixed effect of species, *s*_*ij*_ is 0 for titmouse and 1 for zebra finch, *b*_*0i*_ is the random intercept for each bird, and *∊*_*ij*_ is the residual error. Each full model was compared to a reduced model with the fixed effect of species removed by taking the log-likelihood ratio.

We also assessed whether the difference between species was larger in anterior hippocampus compared to posterior hippocampus. We calculated the average difference in spatial information between species for anterior hippocampus (ΔA) and for posterior hippocampus (ΔP), and compared the difference between these two values (ΔA – ΔP) to the result from a shuffled distribution. The shuffled distribution was generated by randomly assigning cells to either anterior or posterior hippocampus within each bird.

### Analysis of sharp-wave ripples across the long axis

Putative sharp-wave ripples (SWRs) were detected using criteria similar to those used in rodents^14,15^. Some recordings contained sporadic but large movement-related or electrical artifacts; these were detected by setting a threshold for absolute deviation from baseline, chosen separately for each recording, and masking data within 0.4 s of each timepoint above threshold. To detect ripples, the raw LFP was downsampled to 1000 samples/s and then band-pass filtered between 100–200 Hz (Hamming window FIR filter). The analytic signal of the Hilbert transform of the result was calculated (Matlab command *hilbert*), and its magnitude was smoothed with a 4-sigma Gaussian filter to obtain the envelope of ripple power. Putative SWRs were identified as peaks in the ripple envelope that exceeded 5 standard deviations of the mean and stayed above 3 standard deviations for at least 15 ms. The time of each SWR used for alignment in all analyses was the time of the largest ripple trough adjacent to the peak in the ripple envelope. Spectrograms for example single events were generated using a continuous Morse wavelet transformation (Fig. 4a, Extended Data Fig. 8f).

To measure the spatial extent of a single SWR (Fig. 4d), we first detected the number of participating electrodes, as follows. All SWR times were concatenated across all electrodes and sorted, and clusters of inter-event-intervals less than 75 ms were defined as a multi-electrode SWR. The cutoff of 75 ms was determined by inspection of histograms of inter-event-intervals on a log scale, which displayed clear bimodality with a boundary around 75 ms – analogous to methods used to detect bursts of spikes in single neurons^16^. The number of electrodes participating in each multi-electrode SWR was then multiplied by the spacing between electrodes (0.67 mm). Speed and direction of propagation were calculated for events occurring on at least three electrodes by fitting a regression line to the relationship between electrode position and time of the SWR.

To visualize the instantaneous speed and direction of SWRs (**Fig. 4e**), we detected all triplets of SWRs that appeared consecutively on three adjacent electrodes. For each triplet, the correlation coefficient between electrode position and SWR time was calculated. Triplets with a correlation coefficient above 0.9 were deemed propagating triplets, and their velocity was calculated by dividing the spacing between electrodes by the mean inter-event interval:

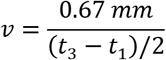

This analysis was repeated separately for SWR events that appeared consecutively on three adjacent electrodes in the posterior→anterior and anterior→posterior directions. Finally, it was repeated for a shuffled dataset in which the times of SWRs for each electrode were circularly shifted by a random time from 0 to the duration of the recording, independent from all other electrodes. This shuffling procedure yielded fewer propagating triplets, and there was no peak in the histogram at faster speeds (Fig. 4e, grey trace).

### Current source density analysis of SWRs

We analyzed the current source density (CSD) during SWRs in three titmice, two of which had electrodes implanted in the transverse plane, and one of which had electrodes implanted in the sagittal plane. All titmice were used to analyze one-dimensional CSD along the radial axis, and the two titmice with electrodes in the transverse plane were additionally used to analyze two-dimensional CSD. For most sessions, the electrodes were advanced in increments of 1/8^th^ of a turn, allowing high-resolution sampling along the radial axis.

To calculate the CSD, the raw LFP from each electrode and each session was first low-pass filtered (40 Hz, Hamming window FIR filter) and averaged across all SWRs. The average voltage at the time of the SWR was then extracted (“sharp wave voltage”). The location of each electrode was determined histologically, creating a map of sharp wave voltages across location in the hippocampus. Only locations with at least 100 detected SWRs were included in the analysis.

For one-dimensional CSD analysis, the position along the radial axis was determined by finding the shortest distance to the lateral ventricle (d_v_) and the shortest distance to the dorsal surface of the hippocampus (d_s_), and then taking the ratio d_v_/(d_v_ + d_s_). For each electrode, the sharp wave voltages recorded across depths were smoothed and upsampled to 101 uniformly spaced positions along the radial axis using a cubic smoothing spline (Matlab command *csaps* with smoothing parameter 0.996). Finally, the one-dimensional CSD was calculated as the negative second spatial derivative of voltage^17^. Only electrodes for which at least 65% of the radial axis was recorded were included. The average CSD across electrodes was then calculated for each bird.

For two-dimensional CSD analysis, we used a cubic spline (smoothing parameter 0.95) to smooth the sharp wave voltages and upsample to 0.01 mm spacing. This map of sharp wave voltages was then spatially differentiated twice to yield the two-dimensional CSD^17^.

### Analysis of theta frequency oscillations

We examined whether theta-frequency oscillations (6−9 Hz)^18^ were over-represented in the hippocampus of birds during a range of behavioral states: movement (periods during the open field and linear track experiments for which the bird was not stationary for more than 5 s), awake immobility (periods during open field and linear track experiments during which the bird was stationary for at least 5 s), and sleep. Power spectra were calculated using Welch’s power spectral density estimate on 1000 separate 1-s segments of the raw LFP for each session. Spectra were averaged across segments, and then averaged across sessions within each bird.

## Notes

### Competing Interest Statement

The authors have declared no competing interest.

